# Factors Affecting Electroconvulsive Therapy Ictal Outcomes: Duration and Postictal Suppression

**DOI:** 10.1101/433789

**Authors:** Wendy Marie Ingram, Cody Weston, Wei Dar Lu, Caleb Hodge, S. Mark Poler, Fatin Nahi, Sharon Larson

**Author notes:** Corresponding author: E-mail address –.

## Abstract

Electroconvulsive therapy (ECT) is an effective and rapid treatment for severe depression, however predictors of therapeutic outcomes remain insufficiently understood. Ictal duration and postictal suppression are two outcomes that may be correlated with patient response, yet patient and treatment variables which may influence these outcomes have not been thoroughly explored. We collected ECT stimulus metrics, EEG parameters, patient demographics, primary diagnosis, and anesthesia type for retrospective ECTs. Univariate and multivariate mixed-effects linear regression models were used to identify variables associated with ictal duration and postictal suppression index. For both outcomes, multivariate models which included all variables resulted in the best fit, reflecting the complex influences of a variety of factors on the ictal response. These results are an important step forward in elucidating patterns in retrospective ECT clinical data which may lead to new clinical knowledge of modifiable factors to optimize ECT treatment outcomes.

## Introduction

Electroconvulsive Therapy, or ECT, is one of the most effective and safe treatments available to psychiatrists for treating severe mood disorders and psychosis. ECT emerged as a therapy due to the observation that schizophrenia seemed to be rare in people with epilepsy. Seizures were originally induced using chemical methods including camphor injection. Later, in 1938, it was found that electrical current could more safely and reliably induce seizures, resulting in improvements in patient symptoms. However, the mechanism of how a controlled seizure leads to reduced depressive symptoms remains poorly understood^1^. Despite popular perception that ECT is antiquated, it can be superior to medication and serves as a vital hope for remission in treatment-resistant cases of depression, bipolar disorder, suicidality and catatonia^2,3^.

Unlike pharmaceutical interventions, ECT requires that practitioners consistently generate a “therapeutic seizure” under continually varying circumstances. For example, a given patient’s seizure threshold is influenced by their age, sex, hydration status, the number of recent seizures that they have had, as well as intrinsic unobservable factors. In turn, the ECT metrics that are involved in stimulating a seizure are numerous. These include percent energy set, stimulus duration, frequency, pulse width, and current. Thus, determining what constitutes “dosage” is not immediately clear in ECT^4^. Consequently, there is considerable variability in the practice of ECT in the United States and around the world.

Optimizing for clinical outcomes requires consideration of the many factors involved in inducing an “adequate” seizure. Early ECT optimization work focused on the prediction of ictal duration, or the duration of the induced seizure^5,6^. More recently it has been determined that ictal duration does not reliably predict therapeutic outcome, but may have utility in limiting cognitive side effects, so long as it is between 25 and 180 seconds^7,8^. Many clinicians continue to optimize for this ictal duration window in their practice today. More recent evidence, however, points to postictal suppression, or the extent to which EEG activity is diminished after the seizure terminates, as being the best predictor of therapeutic outcome^9-12^. Far less is known about which factors predict postictal suppression. As mentioned above, there are many variables that can affect ictal properties and, by extension, clinical outcomes. The identification of individual or combinations of parameters which predict ictal duration and postictal suppression could feasibly improve our ability to reliably produce optimal seizures and thus patient outcomes.

Here we utilize univariate and multivariate linear regression mixed-effects models to determine which patient, stimulus, and EEG parameters predict the continuous outcomes of ictal duration and postictal suppression in a large retrospective study.

## Methods

### ECT Patients and Procedures

All ECT sessions were performed in outpatient surgery at Geisinger Medical Center, located in Danville, Pennsylvania between June 1^st^, 2013 and May 31^st^, 2017. Patients were included in the study retrospectively if they were 18 years or older, had a primary diagnosis of depression (ICD-9 codes: 296.2x, 296.3x, 311; ICD-10 codes: F32.x, F33.x, F43l.x) or bipolar disorder (ICD-9 codes: 296.4x, 296.5x, 296.6x, 296.8x; ICD-10 codes: F30.x, F31.x), and had received an acute series of ECT sessions. Demographic data and ECT parameter summaries are provided in Table 1.

**Table 1.** Demographic Data and ECT parameters. Number and percentages provided. Anesthesia groups are not mutually exclusive.

|  | **Patients**<br>N=84 | **%**<br>- | **ECTs**<br>N=724 | **%**<br>- |
| --- | --- | --- | --- | --- |
| Males | 36 | 43 | 295 | 41 |
| Females | 48 | 57 | 429 | 59 |
| Bipolar | 16 | 19 | 116 | 16 |
| Depression | 68 | 81 | 608 | 84 |
| Pulse width 1.0 | 59 | 70 | 476 | 66 |
| Pulse width 0.5 | 25 | 30 | 248 | 34 |
| Methohexital | 66 | 79 | 555 | 77 |
| Propofol | 28 | 33 | 172 | 24 |
| Ketamine | 12 | 14 | 37 | 5 |

An acute series is defined as 6 or more sequential ECTs, each no more than 3 days apart. ECTs were performed using a Thymatron device (Somatics, LLC. Lake Bluff, IL, USA). Part way through the study period, the machine was changed from a Thymatron DGx with a pulse width of 1.0 mSec to a Thymatron IV with a pulse width of 0.5 mSec. Machine type and pulse width are perfectly correlated variables, thus we included only pulse width in the analysis. The change in machine did not result in any change in what stimulus or ictal measurements were collected.

ECTs were performed in the mornings on Mondays, Wednesdays, and Fridays. Anesthesia and analgesic agents were chosen based on physician experience and preference and included methohexital, propofol, and ketamine which were administered either alone or in combination. For each seizure, the input percent energy set, current, stimulus duration, and frequency were recorded. The ictal outcomes of duration and postictal suppression index were measured and summarized by the Thymatron machines. The procedure start time, the sex and age of the patient, and the number of ECTs in the acute series were also collected from the electronic health record. Bilateral ECT initial dose was based on patient sex (females: 10% energy, 50 mcoul charge; males: 15% energy, 76 mcoul charge) and increased if patients did not respond clinically or if seizures were insufficient during the ECT course (i.e. ictal duration <25 Sec).

We included eleven clinically related features in the model: sex, age, primary diagnosis code, pulse width, anesthesia type, hour of ECT, number of ECT, percent energy set, current, stimulus duration, and frequency. The outcome variables were ictal duration or postictal suppression, both of which are continuous variables. We used mixed effect linear regression to first model the effect of each feature individually on the outcomes of interest. We then created a selective multivariate model through forward selection of only those features that were statistically significant in univariate models. Finally, we created complete multivariate models including all variables and compared model fit criterions described below. Age and sex were included in both selective and complete multivariate models, even if they were not statistically significant predictors in univariate models.

### Statistical analysis

Univariate and multivariate mixed-effects linear regression analyses were performed in R (2017, R Core Team, Vienna, Austria) using package lme4^13^. Visualizations used GraphPad Prism 6 (La Jolla, CA). Patient-specific variability was modeled as a random variable allowing for random patient intercept in all models. Coefficients and restricted maximum likelihood (REML) values for each model are reported and used as model fit criterion. P-values were calculated from associated t-values.

## Results

We evaluated data from 84 patients who underwent a total of 724 ECT sessions. All patients were non-Hispanic white, had median and mean age of 48 years old, ranging from 19 to 88. Patient sex, primary diagnosis, pulse width, and anesthesia numbers and percentages are shown in Table 1 by patient and ECT treatment. Females were slightly more common than males and most patients had a primary diagnosis of depression. A third of ECTs were performed on the second machine, and two thirds of patients received methohexital at least once during an ECT session.

### Ictal duration

Univariate mixed-effects linear regression coefficients and p-values are presented in Table 2. Being administered methohexital was significantly associated with a longer ictal duration while propofol was associated with a shorter duration. The number of ECTs a patient had received in a series, the percent energy, stimulus duration, and frequency were all significantly associated with very minor decreases in ictal duration. The fitness of each mixed-effects model, as estimated by the restricted maximum likelihood (REML) (Figure 1), only changes modestly between each univariate model. Three of the six statistically significant univariate models had the lowest REML values amongst them, indicating the best fit. When these six significant parameters along with patient age and sex are included in a selective multivariate model, the REML value of the model is dramatically reduced, indicating improved fit. Stimulus duration and frequency remain statistically significant, while the other four variables do not. When including all variables in a complete multivariate model, the REML value decreases further, but with only the number of ECTs and the frequency remaining statistically significant predictors of ictal duration.

**Table 2.** Mixed-effects model coefficients and p-values for univariate and multivariate linear regressions of ictal duration and postictal suppression index. Significance: p<0.05*, p<0.01**, p<0.001***. Univar coef = Univariate mixed model coefficients; Selective Multivar coef = multivariate model coefficients including age, sex, and significant univariate variables; Complete Multivar coef = multivariate model coefficients including all univariate model variables.

| Independent variables | Factors | Univar coef | p-value | Selective Multivar coef | p-value | Complete Multivar coef | p-value |
| --- | --- | --- | --- | --- | --- | --- | --- |
| *Ictal Duration* |  |  |  |  |  |  |  |
| Sex | Female (ref) | - | - | - | - | - | - |
|  | Male | -1.70 | 0.63 | 0.95 | 0.77 | 0.65 | 0.84 |
| Age (years) |  | -0.20 | 0.11 | -0.17 | 0.14 | -0.17 | 0.15 |
| Primary dx code | Bipolar (ref) | - | - | - | - | - | - |
|  | Depression | 2.32 | 0.61 | - | - | 4.38 | 0.28 |
| Pulse-width (mSec) | 1.0 (ref) | - | - | - | - | - | - |
|  | 0.5 | -2.99 | 0.34 | - | - | -2.28 | 0.80 |
| Ketamine |  | -3.06 | 0.51 | - | - | 3.21 | 0.58 |
| Methohexital |  | 8.66 | 0.008** | 9.05 | 0.16 | 10.27 | 0.13 |
| Propofol |  | -7.14 | 0.02** | -2.67 | 0.68 | -1.66 | 0.80 |
| Hour of ECT |  | -0.05 | 0.94 | - | - | 0.79 | 0.33 |
| Number of ECTs |  | -0.73 | <0.001*** | -0.37 | 0.06 | -0.39 | 0.05* |
| Percent energy set (%) |  | -0.24 | <0.001*** | 0.08 | 0.53 | 0.08 | 0.59 |
| Current (A) |  | 65.35 | 0.46 | - | - | -89.87 | 0.35 |
| Stimulus duration (Sec) |  | -1.40 | 0.02** | -1.94 | 0.04* | -1.87 | 0.40 |
| Frequency (Hz) |  | -0.35 | <0.001*** | -0.36 | 0.04* | -0.39 | 0.03* |
| *Postictal Suppression Index* |  |  |  |  |  |  |  |
| Sex | Female (ref) | - | - | - | - | - | - |
|  | Male | 0.83 | 0.77 | 1.08 | 0.71 | 0.57 | 0.06 |
| Age (years) |  | -0.07 | 0.48 | -0.11 | 0.28 | -0.13 | 0.18 |
| Primary dx code | Bipolar (ref) | - | - | - | - | - | - |
|  | Depression | 6.23 | 0.09 | 5.82 | 0.11 | 6.62 | 0.06 |
| Pulse-width (mSec) | 1.0 (ref) | - | - | - | - | - | - |
|  | 0.5 | 6.84 | 0.01** | -2.71 | 0.69 | -19.55 | 0.04* |
| Ketamine |  | 15.18 | 0.001*** | 11.17 | 0.02* | 11.64 | 0.02* |
| Methohexital |  | 2.52 | 0.40 | - | - | 0.94 | 0.89 |
| Propofol |  | -3.15 | 0.28 | - | - | -0.13 | 0.98 |
| Hour of ECT |  | -1.83 | 0.03* | -1.45 | 0.09 | -1.60 | 0.06 |
| Number of ECTs |  | -0.15 | 0.39 | - | - | -0.40 | 0.05* |
| Percent energy set (%) |  | 0.02 | 0.63 | - | - | -0.43 | 0.008** |
| Current (A) |  | 7.28 | 0.93 | - | - | 104.85 | 0.31 |
| Stimulus duration (Sec) |  | 1.65 | 0.003** | 1.70 | 0.23 | 7.54 | 0.002** |
| Frequency (Hz) |  | -0.01 | 0.91 | - | - | 0.50 | 0.01** |

**Figure 1.**
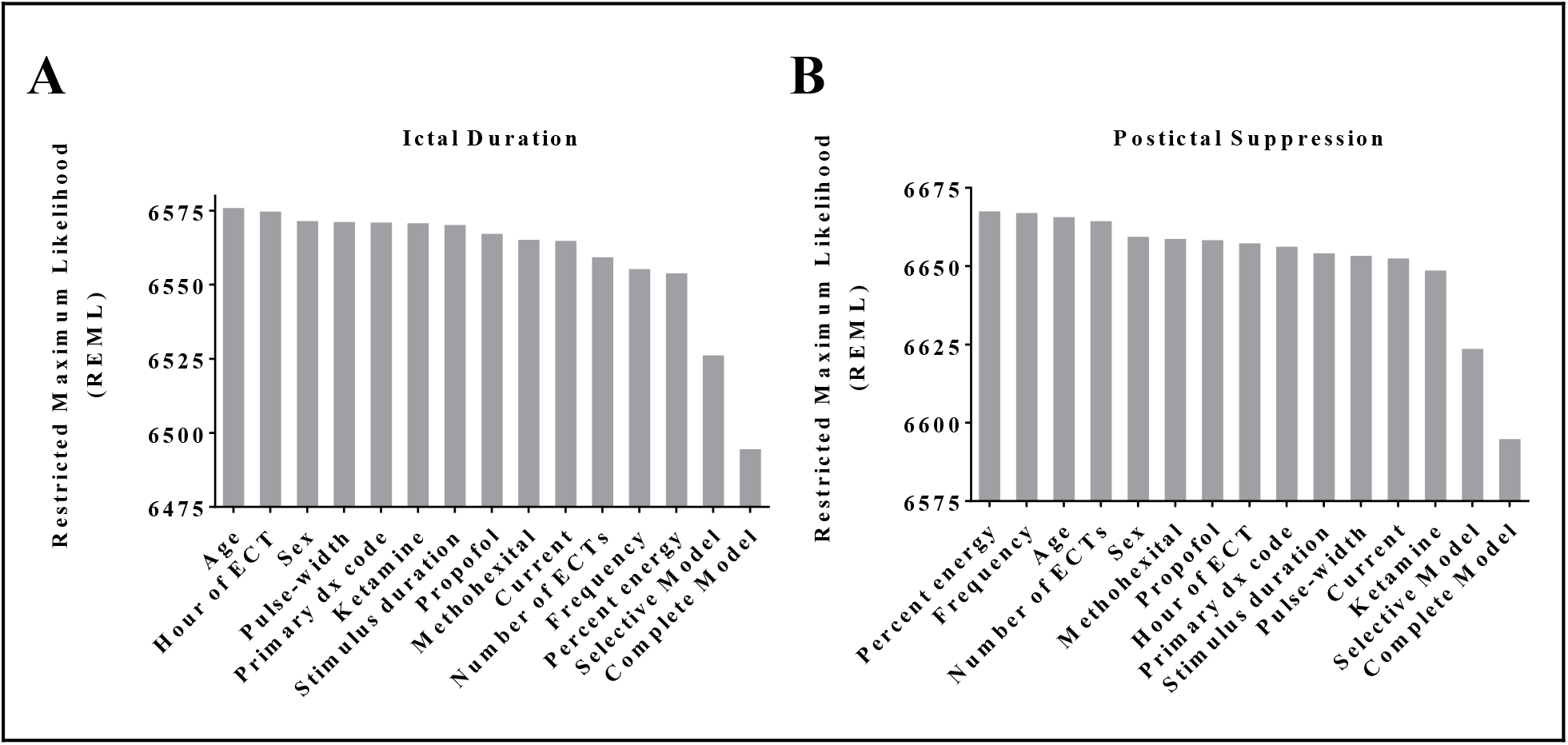
Restricted Maximum Likelihood (REML) of univariate and multivariate mixed-effects models for (A) ictal duration, and (B) postictal suppression, rank ordered. REML is the unbiased estimate of variance and covariance parameters for mixed-effects models, with lower REML relating to better fit of the data.

### Postictal suppression index

Univariate models for postictal suppression index result in pulse width, ketamine administration, and stimulus duration being positively predictive of higher values and statistically significant. These are presented in Table 2. The hour of day during which the ECT is performed is statistically significant and associated with a negative effect on postictal suppression. When these four variables are included with age and sex in a selective multivariate model, only ketamine remains statistically significant with a large positive effect. By including all variables in a complete multivariate model, six variables become statistically significant including those that were significant as univariates (pulse width, ketamine, and stimulus duration) and those that were not (number of ECTs, percent energy, and frequency). Postictal suppression index model REML values were similar in trend to those of ictal duration (Figure 1).

## Discussion

Here we present and compare univariate and multivariate linear mixed-effects models predicting two critical outcome measures of ECT. Ictal duration has been studied extensively due to its previously believed association with therapeutic outcomes and known association with cognitive and other adverse effects. Therefore, its modifiable input parameters have been relatively well studied. Univariate modeling identified expected independent variables predictive of ictal duration. Anesthesia type affected ictal duration, which is consistent with previous studies that show that methohexital is associated with longer seizure time^14-16^. However, we found that the multivariate model that included all independent variables modeled the data best. In this complete model, ECT number and frequency were significant predictors of ictal duration. Frequency is not surprising because it is one of the principle components that can be varied to increase “dosage” and overcome seizure threshold^17,18^. ECT number as a predictor of ictal duration is also consistent with existing literature, as it is well-known that seizure threshold increases with each subsequent ECT administration in an acute series^19^. It will be of interest to explore variable interactions in future mixed-effects modeling and explore the structure of the random effects in this data.

Multivariate modeling of postictal suppression indicates that pulse width, ketamine use, stimulus duration, ECT number, percent energy, and frequency are all significantly predictive of postictal suppression. With the current understanding that postictal suppression is a good predictor of clinical outcome, recognition of these factors has potential to improve clinical practice. The detection of anesthesia choice as a modifier of postictal suppression in this study is consistent with other preliminary studies^15,16^. This is useful information as the standard of care in anesthesia and analgesia for ECT continues to evolve with our understanding of its interplay with clinical outcomes. Previous studies have demonstrated that ketamine may be associated with improved outcomes early in an acute ECT course^20^. The findings of the previous study are provocative since the field currently recommends methohexital as the standard anesthesia method and has not yet produced consistent evidence to recommend ketamine as a primary or adjunct agent^20^. Interestingly, ketamine infusion is emerging as a promising standalone antidepressant treatment^21^, so perhaps ketamine and ECT are interacting in a combinatorial therapeutic way via postictal suppression.

Perhaps the most surprising finding in the present study is that hour of day significantly negatively affects postictal suppression a univariate model. While this variable loses significance in the multivariate models, the size of the effect is considerable and has a near significant p-value of 0.06. Others have recently found that the timing of anesthesia and time of ECT delivery may affect seizure quality but did not look at postictal suppression specifically^22^. This effect may be due to factors including length of time fasting, time since last medication administration, or circadian effects. While the underlying reasons remain unclear and warrant future studies, the recognition of the effect alone might merit the adjustment of patient procedure timing in particularly difficult cases. The key caveat is that it is not clear that time of day was random. It is possible that cases were already stratified according to clinical severity, inpatient/outpatient status, or some other clinical history feature without this information being recorded in the structured EHR.

There are several limitations to this study. The study population was ethnically homogenous and based out of a single center located in rural Pennsylvania. This may limit generalizability to other patient populations nationally and globally. By performing a retrospective study, many factors are not able to be controlled for and confounders could exist due to a lack of randomization. Including all data points in model generation did not allow for cross validation which may result in model overfitting. This can be tested by applying these models to independent datasets in the future. In addition, the data recorded by the ECT machine is generally considered to not be as reliable as expert clinical review of the EEG trace. To estimate the extent of this discordance, future studies may benefit from clinical review of a random sampling to estimate machine/clinician agreement. Another key caveat to be mindful of with clinical interpretation is that the pulse width is correlated perfectly with a change in machine. Thus, it is possible that idiosyncrasies related to the individual machines are being misinterpreted as attributable to pulse width. Future studies at multiple sites as well as larger sample sizes will help to address some of these concerns. In this study we examine the relationship between 11 features and measurable postictal outcomes rather than therapeutic response. This was due to the lack of structured pre- and post-treatment symptom inventory data in the EHR. In future studies, therapeutic improvement may be able to be quantified by extracting clinical assessment of improvement using natural language processing of clinical notes. Additional measures of clinical status such as readmission rates and mortality may assist in estimating therapeutic response. These negative outcomes are rarer, however, and thus larger sample sizes will be required. Additional predictors may be useful to include in future studies, such as current medications, Charlson Comorbidity Index scores, related diagnosis codes, and vital signs.

There are also many strengths of this study. Retrospective studies such as this examine ECT sessions that were performed under clinical conditions on a clinical population, rather than in an artificial setting with selection bias of enrolled participants. The findings presented here are thus immediately relevant to the clinical practice at the investigation site. The unbiased nature of retrospective observational studies can be powerful for hypothesis generation. This repurposing of clinical data is low cost in comparison to randomized controlled trials, thus, there is reasonable cause to thoroughly investigate trends and identify possible predictive factors in existing clinical data prior to conducting prospective studies. In future prospective pragmatic trials these methods may be useful for ongoing analysis and trial arm selection which may reduce the costs and increase the likelihood of adoption by the clinical community. The titration of ECT dosing involves inherent uncertainty and models of ictal outcome may provide a rational basis for guiding care and accelerate the improvement of clinical outcomes. Finally, use of mixed-effects linear regression modeling of many variables both in univariate and multivariate analyses ultimately allows us to begin to unravel the complex interactions and effects that multiple factors have on the ictal outcomes of duration and postictal suppression.

## Conclusions

ECT is just one example of a medical procedure with a multitude of patient, clinician, and treatment mediated variables. For decades, many adjustments of ECT parameters have been driven largely by clinical judgment and preference. In the current era of information technology, it is possible to use historical data to model outcomes of interest and improve care guidelines without having to prospectively randomize patients. The approach demonstrated here could be applied to procedures in many medical domains and represents one high-yield area for informatics techniques to directly improve the health of communities and improve patient centered care.

## Acknowledgements

We would like to thank Jason Brown from the Geisinger Phenomics and Clinical Data Analytics Core for his tireless work refining the data pull, Sarah Robishaw for her help with chart review, and Christopher Bauer, PhD for his excellent comments and suggestions on the manuscript.

